# GuidePro: A multi-source ensemble predictor for prioritizing sgRNAs in CRISPR/Cas9 protein knockouts

**DOI:** 10.1101/2020.07.10.197996

**Authors:** Wei He, Helen Wang, Yanjun Wei, Zhiyun Jiang, Yitao Tang, Yiwen Chen, Han Xu

## Abstract

The efficiency of CRISPR/Cas9-mediated protein knockout is determined by three factors: sequence-specific sgRNA activity, frameshift probability, and the characteristics of targeted amino acids. A number of computational methods have been developed for predicting sgRNA efficiency from different perspectives. We propose GuidePro, a two-layer ensemble predictor that enables the integration of multiple predictive methods and feature sets. GuidePro leverages information from DNA sequences, amino acids, and protein structures, and reduces the impact of dataset-specific biases. Tested on independent datasets, GuidePro demonstrated consistent superior performance in predicting phenotypes caused by protein loss-of-function. GuidePro is implemented as a web application for prioritizing sgRNAs that target protein-coding genes in human, monkey and mouse genomes, available at https://bioinformatics.mdanderson.org/apps/GuidePro.

## Introduction

The CRISPR/Cas9 system has evolved to be the most powerful tool for the perturbation of protein-coding genes, and is widely used in protein functional analysis. The efficiency of CRISPR/Cas9-mediated protein knockout is determined by three factors. First, the activity of an sgRNA impacts the mutation rate at the on-target site. It is now clear that the sgRNA activity is highly dependent on the nucleotide composition of its target DNA, and can be predicted from the sequence [1, 2]. Second, CRISPR/Cas9 introduces small indels to the target DNA sequence that lead to either frameshift or in-frame mutations. While frameshift indels tend to completely abolish protein function, in-frame indels may produce variants that retain the function of the protein[3]. Third, the in-frame indels, which result in the gain or loss of amino acids, may or may not impact protein function, depending on where they occur. A protein is more likely to retain its function with small in-frame indels in nonessential protein regions compared to essential domains [3, 4]. Taken together, these factors collectively contribute to the efficiency of CRISPR/Cas9-mediated protein knockouts.

Corresponding to these three factors, a number of computational methods have been developed for predicting sgRNA efficiency from different perspectives. The majority of the methods are focused on the prediction of sgRNA activity from nucleotide sequence [2, 5–8]. These efforts have been fueled by the development of deep-learning algorithms, which significantly improved predictive power[9–11]. Independent of sgRNA activity, recent studies also showed that the outcomes of CRISPR/Cas9 editing are strongly associated with the target DNA and its surrounding sequences [12, 13]. Machine learning approaches have enabled the prediction of indel types and the frameshift/in-frame probability at the Cas9 cutting site, based on the local nucleotide sequence [13–15]. Moreover, we and others have shown that the “importance” of targeted amino acids are predictable from conservation, secondary structure, and post-translational modifications [16, 17], which allow the assessment of targeting efficiency at a protein level. Therefore, it is plausible to integrate sgRNA activity, indel types and amino acid information to further improve sgRNA efficiency prediction.

To take advantage of existing predictive methods by leveraging the information from their outputs, we developed GuidePro, a two-layer ensemble predictor that enables the integration of multiple predictive methods and feature sets. Tested on independent datasets, GuidePro demonstrated consistent superior performance in predicting phenotypes caused by protein loss-of-function, suggesting its robustness in a broad spectrum of experimental settings.

## Results and Discussion

A schematic overview of the GuidePro framework is shown in Fig.1a. We model the knockout efficiency, *E*, to be a function dependent on the three factors: sgRNA activity (SA), frameshift probability (FP), and amino acid sensitivity (AS):

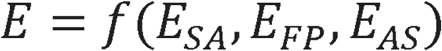

**Fig. 1.**
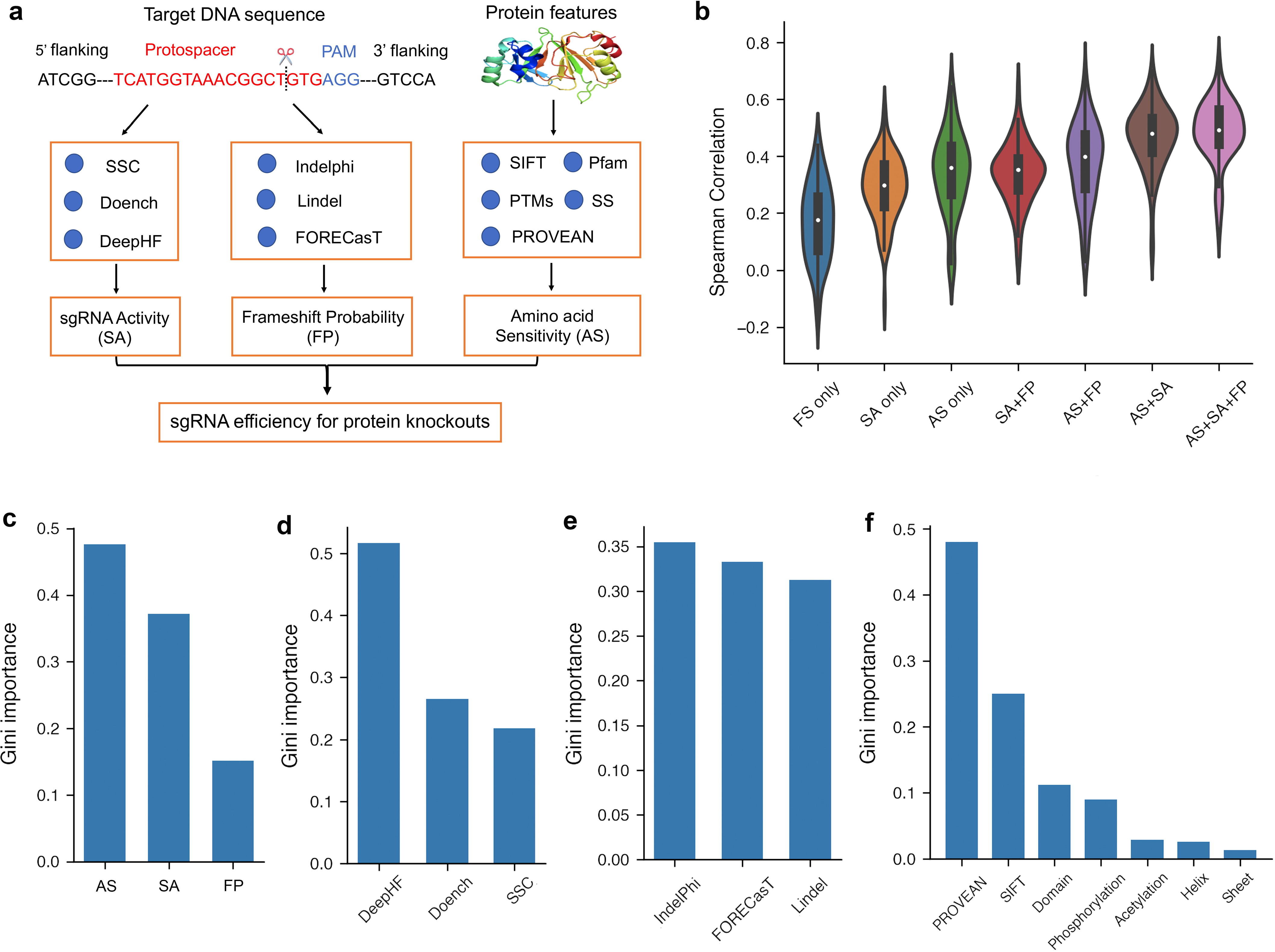
GuidePro integrates multiple factors for predicting protein knockout efficiency with CRISPR/Cas9. **(a)** A schematic view of GuidePro framework, which consists of two layers: the first layer estimates three factors that jointly contribute to protein knockouts, including sgRNA activity (SA), frameshift probability (FP) and amino acid sensitivity (AS). The second layer combines three factors to predict the final sgRNA efficiency for protein knockouts. **(b)** Violin plots comparing the performance of individual factors and their combinations using Spearman correlation coefficient between the predicted and measured knockout effects through leave-one-gene-out cross validation. **(c)** The feature importance of the three factors in the combined predictive model, measured by Gini importance scores. **(d-f)** The Gini importance of the features in the estimation of individual factors, including **(d)** sgRNA activity (SA), **(e)** Frameshift probability (FP), and **(f)** Amino acid sensitivity (AS).

The term *E_SA_* represents the sgRNA activity that contributes to knockout efficiency. To leverage the power of multiple predictors, we model *E_SA_* to be a non-linear combination of the outputs of three methods, DeepHF[10], Doench method[8], and SSC[2], which were previously trained on independent datasets of various experimental readouts. Similarly, we model the frameshift probability *E_FP_* to be a function of the outputs of inDelphi [13], Lindel [14], and FORECasT[15], which were developed for indel type prediction. The amino acid sensitivity *E_AS_* is dependent on the predictions and annotations of protein features, including conservation (PROVEAN and SIFT scores), domain annotations, post-translational modifications (PTMs) and secondary structures [18–22]. Taken together, the GuidePro framework is organized into a two-layer assembly structure, in which the first layer estimates three individual factors and the second layer combines the factors into a final score.

The main goal of GuidePro is to prioritize all sgRNAs that target the coding exons of a protein for efficient knockout. To train the model in an unbiased sample space, we selected the Munoz data, a large tiling-sgRNA dataset that includes all possible sgRNAs targeting exons of over 100 genes in three cell lines[4]. In this dataset, the sgRNA efficiency is measured to be the dropout z-score in cell viability screens. Our preprocessing step left 25,079 sgRNAs targeting 91 genes for the analysis (Table S1). The sgRNAs were randomly split half-and-half for the training of the first and the second layers, respectively. We used a bootstrapping strategy to minimize the variation caused by random sampling (see Supplementary method).

To test if an integration of the three factors could better explain the variations of sgRNA efficiency, we computed the Spearman correlations of predicted and observed efficiency measures in settings of individual factors or their combinations (Fig. 1b). As expected, a combination of all three achieved the highest average correlation of 0.523 over the 91 genes. Of note, different machine learning methods (Random Forester and SVM) achieve similar performance (Fig. S1). Feature importance analyses using multiple approaches indicate that the amino acid sensitivity and sgRNA activity are the two major determinants of knockout efficiency, whereas the frameshift probability contributes to the efficiency to a lesser degree (Fig. 1c and Fig. S2a, 1e). In the estimation of sgRNA activity, DeepHF is the most important feature compared to the others, suggesting a significant improvement made by the deep-learning algorithm in DeepHF (Fig. 1d and Fig. S2b, 1f). The three indel type prediction methods almost equally contribute to the estimation of frameshift probability (Fig. 1e and Fig. S2c, 1g). Consistent with previous findings[16, 17], conservation scores predicted by PROVEAN or SIFT demonstrate much greater importance than other protein features in the estimation of amino acid sensitivity (Fig. 1f and Fig. S2d, 1h).

Next, we asked if GuidePro could improve the design of the sgRNA libraries for high-throughput functional screens. We collected 13 screening datasets [17, 23–26] and compared the performance of GuidePro with 8 existing tools based on the correlation of predicted efficiency and cell phenotypes (Tables S2-4). These datasets were divided into two groups. The first group includes the data that were used to train the methods, which are considered “dependent”. The second group consists of “independent” datasets that are ideal for an unbiased evaluation. In the dependent group, most existing methods are subject to overfitting; that is, a method achieves high correlation on training datasets but the performance degrades when the method is applied to other datasets (Fig. 2a). The overfitting problem is likely due to the variations in protocols, vector systems, and library design strategy in CRISPR/Cas9 screens. Since GuidePro combines the outputs of several methods, it is less sensitive to overfitting and performs consistently well. On the independent group of datasets, especially on tiling-sgRNA data, GuidePro significantly outperforms other methods in predicting phenotypes (Fig 2b), suggesting its robustness for the design of sgRNA libraries in various applications of CRISPR/Cas9 screens.

**Fig. 2.**
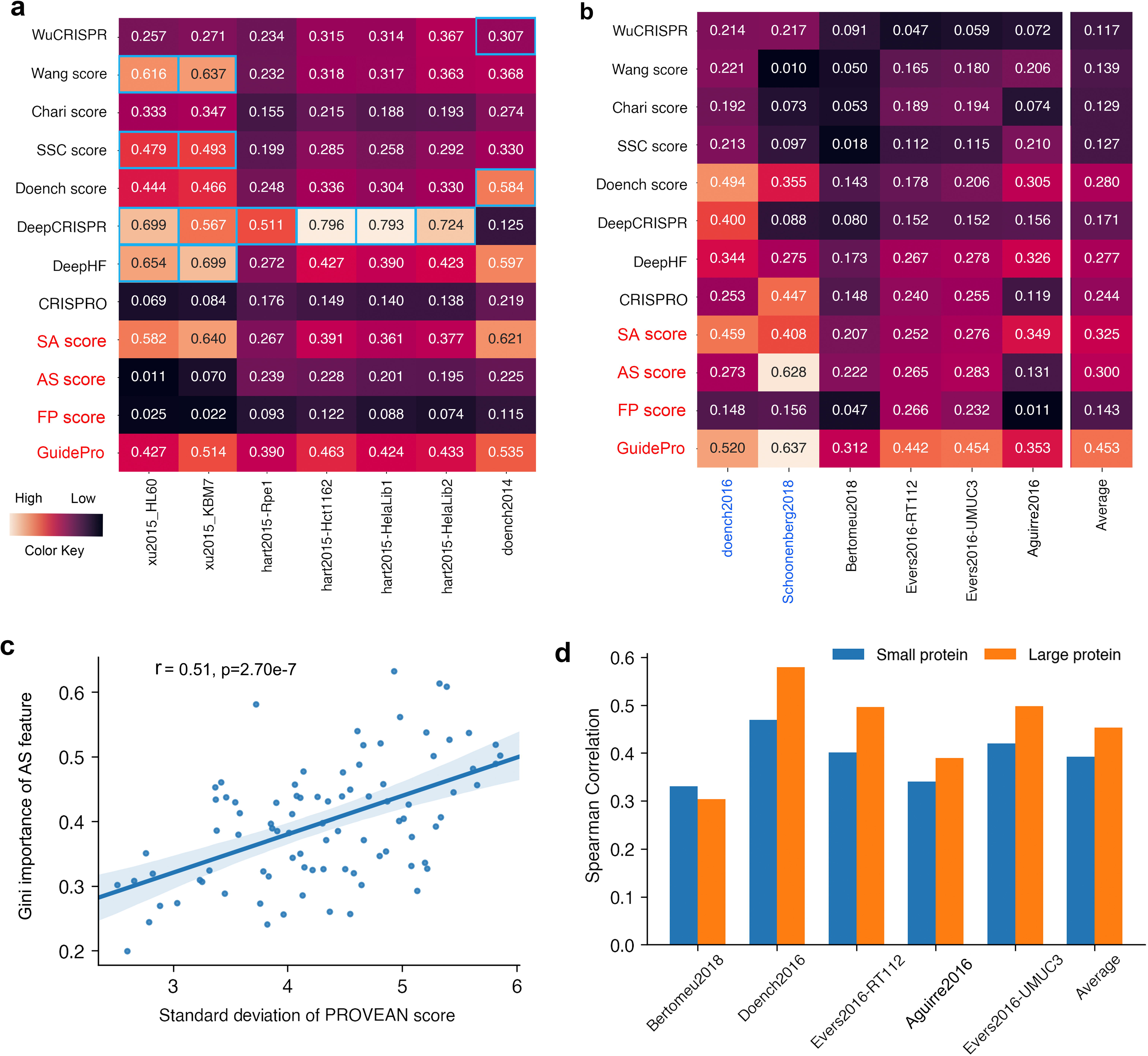
An evaluation of GuidePro. **(a, b)** Heatmaps of Spearman correlation coefficients between predicted and measured knockout efficiency for GuidePro and existing computational methods on **(a)** dependent datasets and (**b)** independent datasets. Three individual factors (SA, AS, FP) and the combined model (GuidePro) are marked in red. The datasets with tiling-sgRNA design are marked in blue. The correlation of methods corresponding to their own training datasets are highlighted with blue rectangular frames. **(c).** Scatter plots showing the correlation of the standard deviations of PROVEAN score against the Gini importance of AS features. The p-value was calculated with Pearson correlation coefficient test. **(d).** Bar plots comparing the performance of GuidePro for sgRNAs targeting large proteins (>500 AA) versus small proteins (<500 AA) on independent datasets (p=0.032, paired t-test).

In the Munoz data, the amino acid sensitivity (AS) is the most important factor governing sgRNA efficiency (Fig. 1c). We examined if the AS features also significantly contribute to sgRNA efficiency in other datasets. Interestingly, the AS features predict phenotypes to various degrees in a library-dependent manner (Fig. 2a and 2b). There are two explanations for this observation. First, the sgRNA sample spaces vary among different libraries. Some libraries were optimized to target amino acids that are more likely to be functional, evidenced by the fact that the sgRNAs in these libraries are associated with highly conserved amino acids compared to randomly selected sgRNAs (Fig. S3). Therefore, only a smaller fraction of variations of phenotypes in these datasets is explainable by the AS features. Second, the contribution of AS features is also protein-dependent. If a vast majority of amino acids in a protein are indispensable, the variation of sgRNA efficiency will be dominated by sgRNA activity rather than amino acids. Indeed, we found that the importance of AS in sgRNA efficiency prediction is positively correlated with the variation of amino acid conservation (Fig. 2c, Fig. S4), suggesting that the AS features are of higher importance for large, multi-domain proteins that underwent diverse evolution processes. Consistently, GuidePro is associated with higher predictive power for larger proteins (>500 AA) compared to that for smaller proteins (<500 AA) (Fig. 2d).

To facilitate users in selecting optimal sgRNAs for efficient protein knockout, GuidePro is implemented in a web application that includes exome-wide prediction in human, monkey and mouse genomes, available at https://bioinformatics.mdanderson.org/apps/GuidePro.

## Availability of data and materials

Detailed methods and other processed data are available in **supplementary data** in this article, and source codes for GuidePro model training and testing are available at https://github.com/MDhewei/GuidePro. All other relevant data can be obtained from the authors upon request.

## Acknowledgements

This work was supported by CPRIT grant RR160097 (H.X.). H.X is a CPRIT scholar in cancer research.

## Contributions

W.H. and H.X. conceptualized the study. W.H. developed the method and web application with the help from H.W., Z.J., Y.W. and Y.C., W.H., Y.T. and H.W. collected the datasets and performed computational analysis. H.X. supervised the project. All authors participant in writing the manuscript.

