## Supplementary method for "GuidePro: A multi-source ensemble predictor for prioritizing sgRNAs in CRISPR/Cas9 protein knockouts"

**Supplementary Methods**

1. **Description of the GuidePro framework**

GuidePro integrates three factors that jointly contribute to protein knockouts: i) sgRNA activity (SA), which adopts the output scores of three sequence-based prediction tools as input features. ii) Frameshift probability (FP), which combines the indel type prediction of three different machine learning models. iii) Amino acid sensitivity (AS), which is predicted from protein features derived from public databases and other prediction tools.

**1.1 sgRNA activity**

The output scores of three different computational tools used for sgRNA on-target activity prediction are adopted as input features: SSC score [1], Doench score [2] and DeepHF score [3]. SSC score is calculated with our own source code. Doench score is calculated with Azimuth 2.0 which is available from <https://github.com/MicrosoftResearch/Azimuth>. DeepHF source code is downloaded from <https://github.com/izhangcd/DeepHF>.

**1.2 Frameshift probability**

The results of three machine learning models predicting CRISPR knockouts outcomes are used to predict indel types: Indelphi [4], FORECasT [5] and Lindel [6], whose source codes are publicly available from <https://github.com/maxwshen/inDelphi-model>, <https://github.com/felicityallen/SelfTarget> and <https://github.com/shendurelab/Lindel>, respectively.

**1.3 Amino acid sensitivity**

We derived protein features that are associated with amino acid sensitivity to CRISPR knockouts from publicly available protein annotation databases and prediction tools: domain annotation [7], conservation [8, 9], post-translational modifications (PTMs) distribution [10] and secondary structure [11]. Then we transformed them to different mathematical expressions that best describe these features:

Given a protein with N amino acids: (A_1_, A_2_, A_3_, A_4_, …, A_n_), the domain annotation is defined as:

$$Domain\left( A_{i} \right)=\left\{ \begin{aligned} 1, &if A_{i}inside Pfam domain \\ 0, &if A_{i}outside Pfam domain \end{aligned} \right.$$

Protein secondary structure annotation is defined as:

$$SS\left( A_{i} \right)=\left\{ \begin{aligned} \left[ 1,0 \right], if predicted as helix \\ \left[ 0,1 \right], if predicted as strand \\ \left[ 0,0 \right], if predicted as coil \end{aligned} \right.$$

For protein conservation, we first calculate SIFT score and PROVEAN score for each amino acid as S = (S_1_, S_2_, S_3_, S_4_, …, S_n_) and P = (P_1_, P_2_, P_3_, P_4_, …, P_n_). Considering that conservation scores of different amino acids are not independent but associated with each other, we smoothed the scores at each amino acid using Kernel Density Estimation as follows:

$S(A_{i}$*) =* $\sum_{i=1}^{n} K(\frac{S_{i}-S}{h})$* $\sum_{i=1}^{n} S_{i}$

$P(A_{i}$*) =* $\sum_{i=1}^{n} K(\frac{P_{i}-P}{h})$* $\sum_{i=1}^{n} S_{i}$

$A_{i}$ is the i-th amino acid of a certain protein, $S_{i}$ denotes the corresponding SIFT score of $A_{i}$, and $P_{i}$ denotes the corresponding PROVEAN score of $A_{i}$. K is the Gaussian kernel and h is the bandwidth parameter.

For PTMs distribution, let M = (M_1_, M_2_, M_3_, M_4_, …, M_n_) represent certain types of PTM at each amino acid:

$$M_{i}=\left\{ \begin{aligned} 1, &if A_{i} harbors certain PTM \\ 0, &if no certain PTM at A_{i} \end{aligned} \right.$$

Considering that PTM at a certain amino acid may also affect its neighboring amino acids, we smoothed the PTM value at each amino acid using Kernel Density Estimation:

$M(A_{i}$*) =* $\sum_{i=1}^{n} K(\frac{M_{i}-M}{h})$ * $\sum_{i=1}^{n} M_{i}$

Where $A_{i}$ is the i-th amino acid of a certain protein and $M_{i}$ denotes the corresponding PTM value at $A_{i}$. K is the Gaussian kernel and h is the bandwidth parameter. We reasoned that two amino acids that are 10 AA away may have little association with each other, so we generalized the bandwidth of all the kernel density estimation models to 10.

2. **Model Training**
   2.1 **Preprocessing of training set**

A publicly available tiling CRISPR screen with all possible sgRNAs targeting exons of 153 protein-coding genes was utilized as the training set [12]. The dataset includes tiling-sgRNA screens on three cell lines (RKO, NCI-H1299, and DLD-1). We first computed the average Z-score for each gene in each cell line. Using a threshold of -0.4, as suggested in the original publication, we identified 80, 87, and 90 essential genes for RKO, NCI-H1299, and DLD-1 cell lines, respectively. Only genes that are essential in at least one cell line were kept which resulted in 91 genes with 25079 sgRNAs left in the final training set. For each gene, we averaged the Z-scores in the corresponding essential cell line(s) for each sgRNA to increase the signal-noise ratio, then calculated the mean-centered Z-score as the final measurement for the knockout efficiency.

**2.2 Models used for training**

Two commonly used machine learning models were utilized for the training of GuidePro: a support vector regression model with RBF kernel (SVM-RBF) and a random forest regression model (RF), which were implemented through SVR and RandomForestRegressor modules from the scikit-learn package in python [13], respectively. The optimized parameters for SVM-RBF and RF were obtained using GridSearchCV from the scikit-learn package.

We tried different combinations of two machine learning models to train two layers of GuidePro: RF-RF, RF-SVM, SVM-RF and SVM-SVM. The sgRNAs were randomly split half-and-half for the training of the first and the second layers, respectively. We used a bootstrapping strategy to minimize the variation caused by random sampling. To evaluate the performance of different model combinations, we adopted a cross validation strategy by leaving out sgRNAs targeting one gene at a time (leave-one-gene-out). Spearman correlation coefficient between predicted and measured knockout efficiency was used for the evaluation. We found similar performance of different model combinations (Fig. S4). We selected SVM-SVM as our final prediction model, which obtained relative better performance (averaged Spearman’s correlation coefficients: 0.523).

**2.3 Feature importance**

We used three different ways to measure the feature importance: i). Gini importance, which refers to the decrease in the mean-squared error (the criterion used to train each regression tree) when that feature is introduced as a node in the tree. The Gini importance was implemented through the RandomForestRegressor module from the scikit-learn package in python. ii). Permutation importance, which is defined to be the decrease in a model score when a single feature value is randomly shuffled. iii). Drop-column importance, which is measured by the difference between the baseline importance of a feature calculated through permutation and the score from the model missing that feature. The permutation importance and drop-column importance were implemented through the permutation_importances module and the dropcol_importances module from the rfpimp package, respectively. The rfpimp packages can be obtained from <https://github.com/parrt/random-forest-importances>.

3. **Comparing with other methods and tools**

We collected 13 different CRISPR screen datasets to compare the performance of GuidePro to other existing tools, among which 7 datasets were previously used for the training of different methods (dependent datasets) and 6 datasets haven’t been used for the training of any methods involved in the comparison (independent datasets). The dependent datasets are derived from benchmarking datasets collected by Haeussler et al.[14].The independent datasets incudes a tiling CRISPR screen targeting two key erythroid transcription factors [15], a CRISPR knockout screen focused on 43 essential genes [16], and three genome-wide CRISPR screens [17, 18]. In the genome-wide screen data, only sgRNAs targeting the pan-essential genes which are identified in the previous publication were kept for testing [19].

The performance of each prediction model was evaluated by Spearman’s correlation coefficients between experimentally measured sgRNA activities and prediction scores from each model. The correlations of models tested against their own training datasets were excluded for fair comparison.

4. **Exome-wide prediction of sgRNA efficiency for protein knockouts**

Based on our new sgRNA knockout efficiency prediction approach, we performed exome-wide prediction of sgRNA efficiency for protein knockouts in human, monkey, and mouse genomes with the following steps: Step 1: Extract genomic sequence of all the exons for the target gene. The transcript annotation of exons was downloaded from the NCBI CCDS database [20]. Step 2: Search for all the possible sgRNAs targeting exons of the target gene with an NGG PAM sequence. Annotate the target genomic loci and the corresponding amino acid position on the protein. Step 3: Annotate the protein features, sgRNA sequence features, and frameshift probabilities for all the sgRNAs as described in section 1. Step 4: Predict protein knockout efficiency with GuidePro and prioritize the sgRNAs with high predicted efficiency.

We provide a user-friendly web tool for selecting the optimized sgRNAs with high efficiency for the knockout of a specific protein-coding gene in human, monkey, and mouse genomes at <https://bioinformatics.mdanderson.org/apps/GuidePro>. Users can simply select the genome and gene name of interest and the server will output all the sgRNAs targeting the exons of the gene ranking from high predicted efficiency to low.
