## Supplementary Figures for "GuidePro: A multi-source ensemble predictor for prioritizing sgRNAs in CRISPR/Cas9 protein knockouts"

He et al.

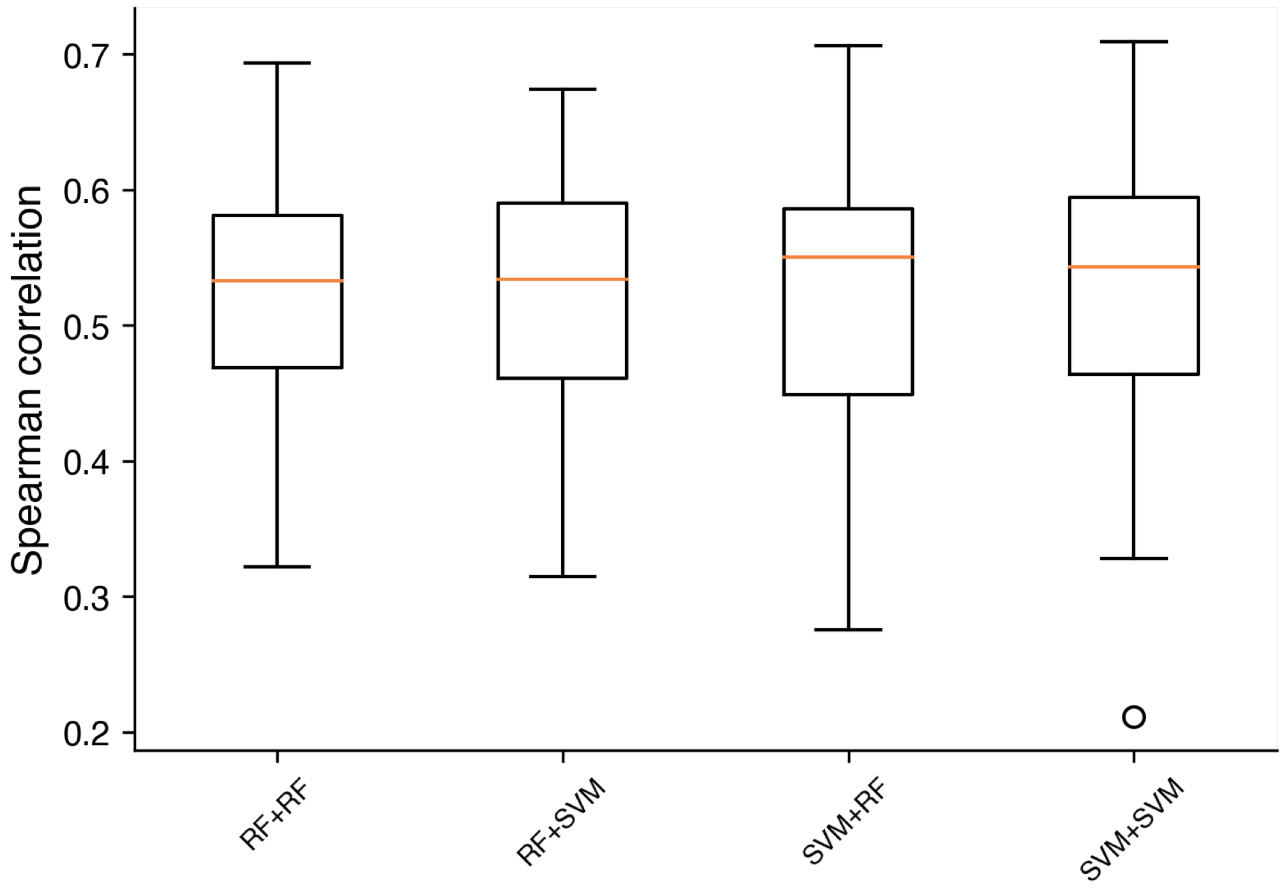


**Figure S1**. Box plots comparing the performance of different model combinations using leave-one-gene-out cross validation.

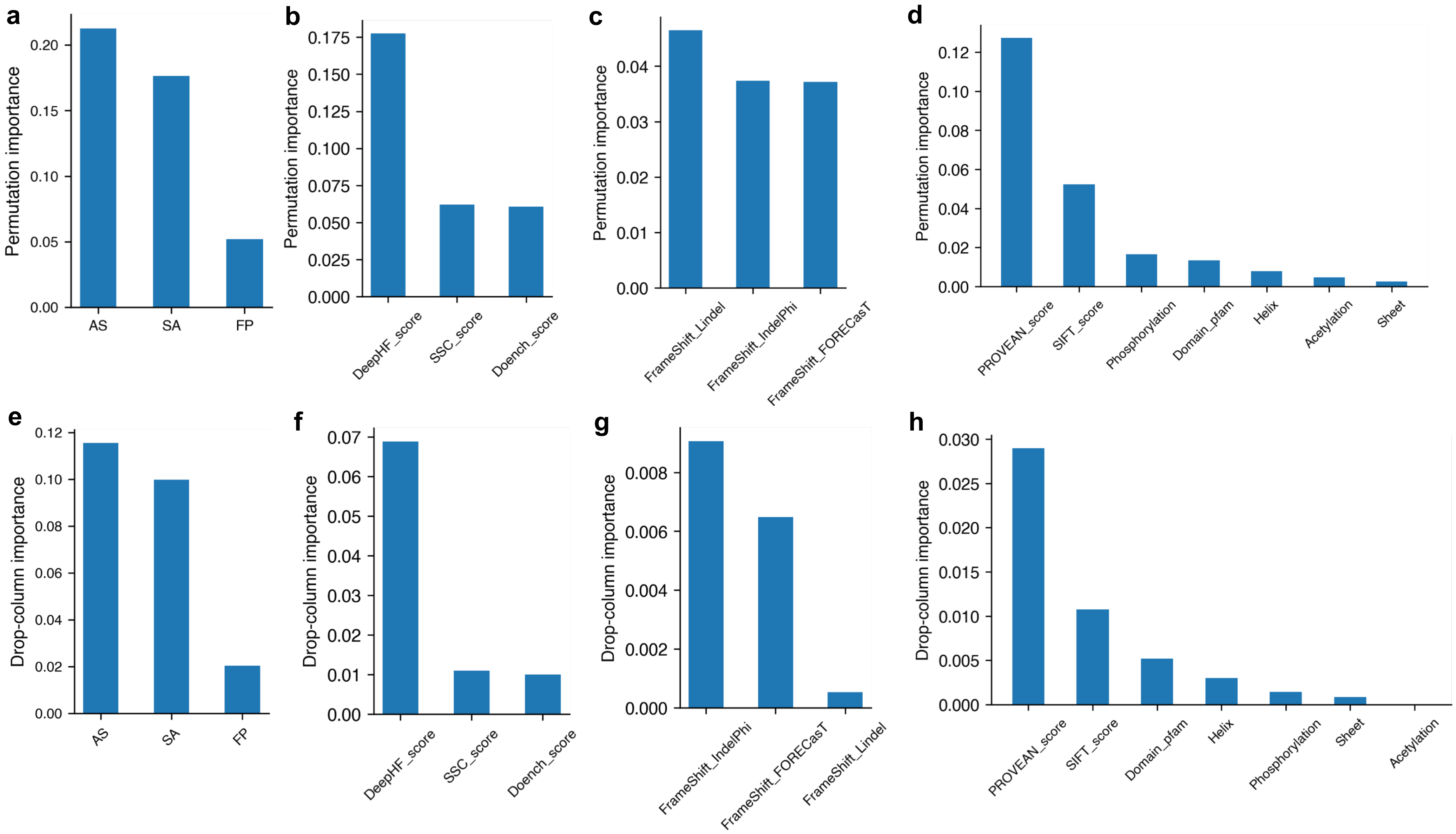


**Figure S2**. Permutation importance and drop-column importance for features in **(a) and (e).** The combined prediction model (GuidePro); **(b) and (f).** sgRNA activity (SA) predictor; **(c) and (g).** Frameshift probability (FP) predictor; **(d) and (h)**. Amino acid sensitivity (AS) predictor.

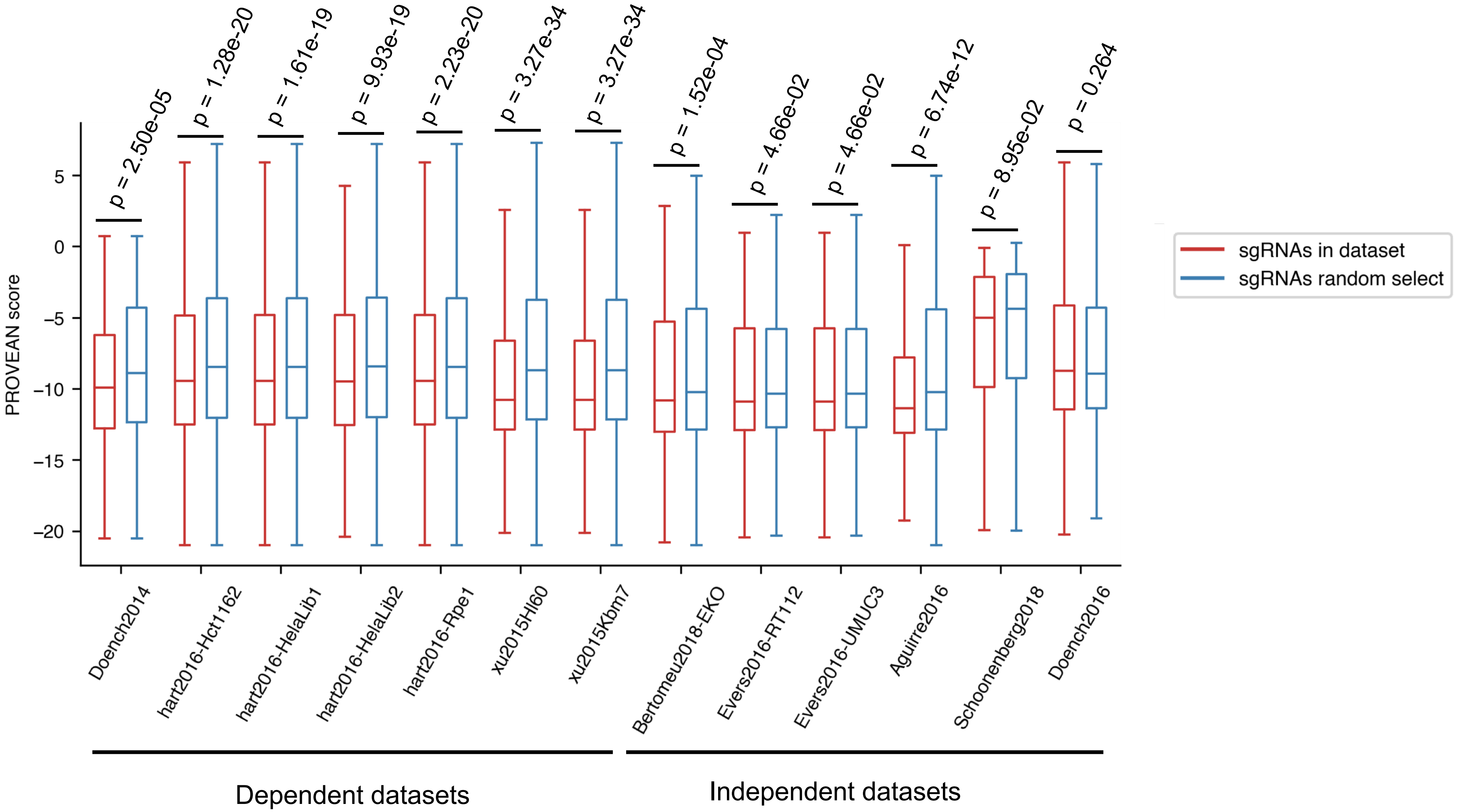


**Figure S3**. Box plots comparing the PROVEAN score of amino acids targeted by sgRNAs in the testing datasets versus sgRNAs randomly selected from tiling-sgRNAs targeting same gene. The p-values were calculated using Mann-Whitney test.

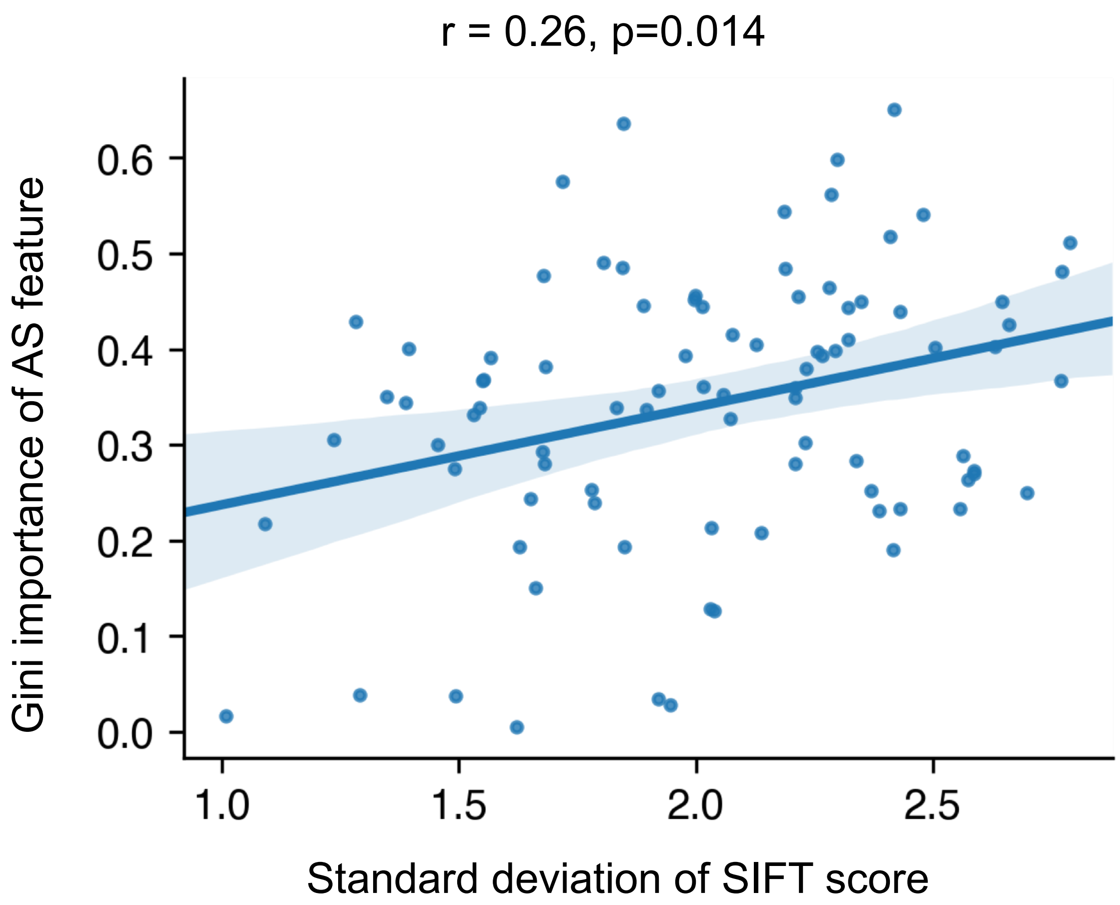


**Figure S4**. Scatter plot showing the correlation between standard deviation of SIFT score and Gini importance of AS features for knockout efficiency prediction. The p-value was calculated with Pearson correlation coefficient test.
